# Contrasting effects of reduced phosphatidylcholine synthesis in absence of seipin: embryonic lethality goes down while lipid droplets get even larger

**DOI:** 10.1101/2022.04.10.487805

**Authors:** Jinglin Zhu, Sin Man Lam, Leilei Yang, Jingjing Liang, Mei Ding, Guanghou Shui, Xun Huang

**Author notes:** To whom correspondence should be addressed at: State Key Laboratory of Molecular Developmental Biology, Institute of Genetics and Developmental Biology, Chinese Academy of Sciences, Beijing 100101, China, 86-10-64806560 (Tel&Fax).

## Abstract

Seipin plays a vital role in lipid droplet homeostasis and its deficiency causes congenital generalized lipodystrophy type II in humans. It is not known whether the physiological defects are all caused by cellular lipid droplet defects. Loss-of-function mutation of *seip-1*, the *C. elegans* seipin ortholog, causes embryonic lethality and lipid droplet abnormality. We uncover *nhr-114* and *spin-4* as two suppressors of *seip-1* embryonic lethality. Mechanistically, *nhr-114* and *spin-4* act in the “B12-one-carbon cycle-phosphatidylcholine (PC)” axis and reducing PC synthesis suppresses the embryonic lethality of *seip-1* mutants. Conversely, PC deficiency enhances the lipid droplet abnormality of *seip-1* mutants. The suppression of *seip-1* embryonic lethality by PC reduction requires polyunsaturated fatty acid (PUFA). Therefore, seipin and phosphatidylcholine exhibit opposite actions in embryogenesis, while they function similarly in lipid droplet homeostasis. Our results demonstrate that seipin-mediated embryogenesis is independent of lipid droplet homeostasis.

**Highlights:**

1. *seip-1* suppressors act in the “B12-one-carbon cycle-PC” pathway.
2. Reducing PC synthesis suppresses the embryonic lethality of *seip-1* mutants.
3. Suppression of the embryonic lethality by PC reduction requires PUFA.
4. Reduced PC synthesis enhances the large lipid droplet of *seip-1* mutants.

## Introduction

Lipids are essential for life and take part in nearly all physiological processes. Abnormal lipid metabolism is associated with many diseases, including developmental, neuronal, and reproductive diseases, as well as metabolic syndromes. Regulation at both the cellular and tissue levels is required to maintain the organismal homeostasis of lipid metabolism. At the cellular level, lipid droplets, which originate from the endoplasmic reticulum (ER), are hub organelles for neutral lipid storage and utilization (Walther and Farese, 2012; Welte, 2015).

Seipin, an integral ER protein, plays an important role in lipid droplet homeostasis (Fei et al., 2008; Szymanski et al., 2007). Located at the contact site between the ER and the budding nascent lipid droplet, seipin possesses a luminal lipid-binding motif between its two transmembrane domains and forms oligomers (Sui et al., 2018; Yan et al., 2018). Seipin stabilizes nascent lipid droplets and promotes their growth by facilitating the transfer of neutral lipid from the ER into the associated lipid droplets. In seipin-deficient cells, abnormal partitioning of neutral lipids results in the formation of tiny lipid droplets and some supersized lipid droplets (Salo et al., 2019; Wang et al., 2016). Besides seipin, other proteins or lipid factors, including FIT, Snx14, ACSL3 and phosphatidylcholine (PC), have also been identified to promote lipid droplet biogenesis and/or lipid droplet growth (Choudhary et al., 2015; Datta et al., 2019; Fujimoto et al., 2007; Gao et al., 2019; Krahmer et al., 2011).

In addition to the cellular lipid droplet defect, seipin deficiency also causes Bernardinelli-Seip congenital lipodystrophy 2 (BSCL2)/congenital generalized lipodystrophy type II (CGL2) in humans (Magre et al., 2001). The patients lose nearly all their subcutaneous fat tissue and develop many metabolic syndromes, including fatty liver, diabetes and hypertriglyceridemia, as well as many non-metabolic disorders, such as muscular hypertrophy, mental retardation and sperm abnormality (Gomes et al., 2009; Jiang et al., 2014). It is not known how seipin deficiency causes so many physiological defects. In particular, it is unclear whether all these defects are due to abnormal lipid droplet homeostasis.

Seipin is conserved from yeast to human. Similar to BSCL2, numerous physiological defects have been reported in *seipin*-deficient animal models (Bai et al., 2020; Cui et al., 2011; Ebihara et al., 2015; Jiang et al., 2014; Zhou et al., 2014). *seip-1*, the only ortholog of human seipin in *C. elegans*, regulates the homeostasis of intestinal lipid droplets and also embryogenesis. Here, through the identification and characterization of *seip-1* suppressors, we found that reducing PC synthesis enhances the lipid droplet defect of *seip-1* mutants, while suppresses the embryonic lethality. The suppression of *seip-1* embryonic lethality by PC reduction requires polyunsaturated fatty acid (PUFA). Therefore, this suggests that seipin-mediated embryogenesis is independent of lipid droplet homeostasis.

## Results

### Deletion of *seip-1* results in embryonic lethality

*tm4221*, a deletion allele of *seip-1* with a 299-bp deletion including exon 4 and part of exon 3, was obtained from the *C. elegans* Gene Knockout Consortium (Figure 1A). We found that *seip-1(tm4221)* mutants exhibited highly penetrant embryonic lethality, whereas brood size was not significantly changed (Figure 1B, C, D). An extrachromosomal array expressing the wild-type *seip-1* gene partially rescued the embryonic lethality of *seip-1(tm4221)* mutants (Figure 1E). These observations suggest that SEIP-1 is critical for *C. elegans* embryonic development.

**Fig. 1.**
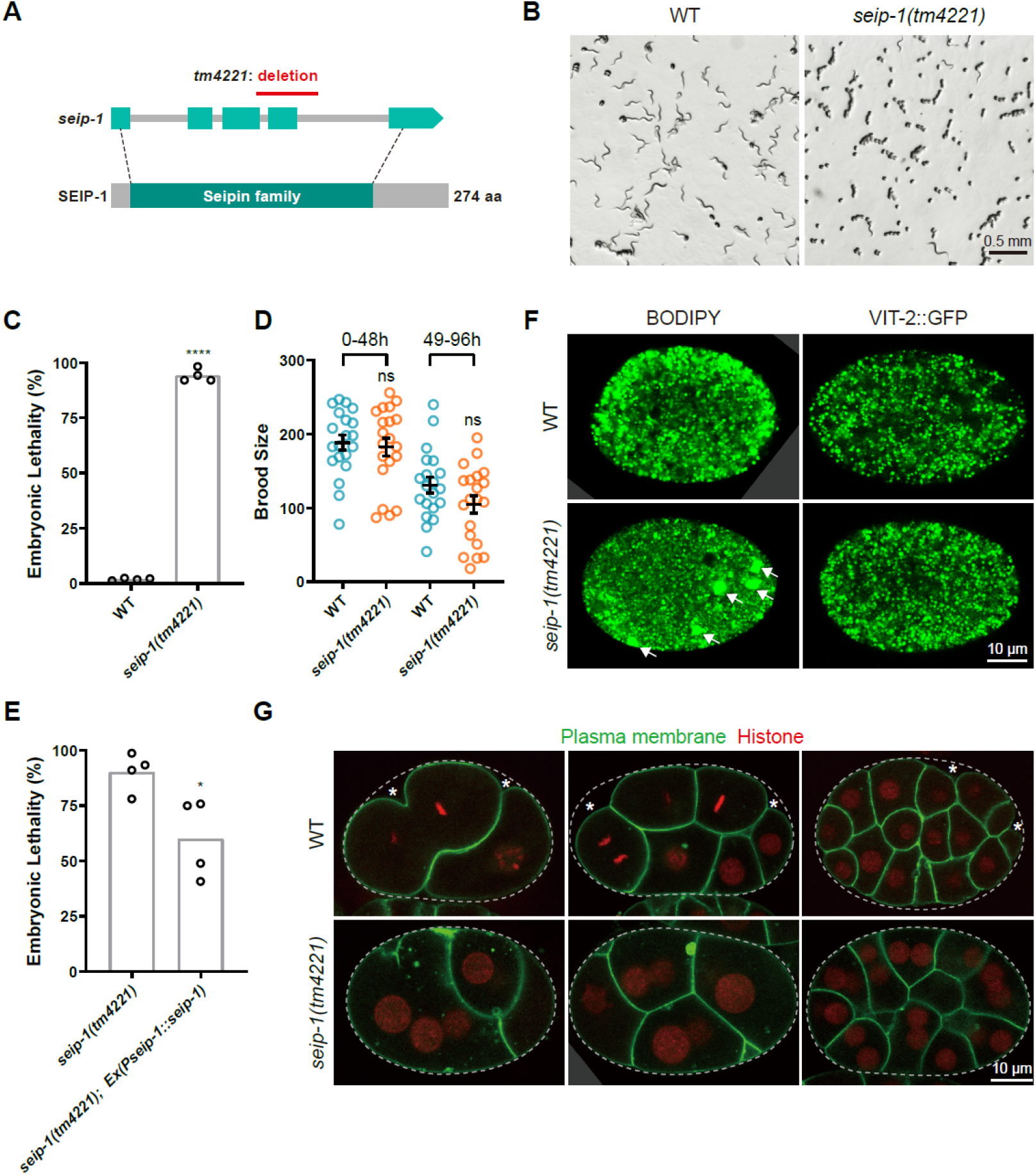
*seip-1(tm4221)* mutants exhibit embryonic lethality and have large lipid droplets. (A) Diagram of the open reading frame and the conserved protein domain of *C. elegans* seipin. Light cyan rectangles indicate the exons and grey lines indicate the introns in the *seip-1* gene. The dark cyan rectangle highlights the conserved region in the SEIP-1 protein. Dashed lines indicate the start and end of the region that encodes the conserved protein domain. The red line indicates the deleted genomic region in the *seip-1(tm4221)* mutant. (B) Bright-field images of worm plates containing wild type and *seip-1(tm4221)* mutant animals. There are many dead embryos and few larvae on the *seip-1(tm4221)* mutant plate. (C) Quantification of embryonic lethality of wild type and *seip-1(tm4221)* mutants. Each point represents a biological repeat. Statistical significance was determined by two-tailed unpaired t-test with Welch’s correction: \*\*\*\**P* < 0.0001. (D) Quantification of brood size of wild type and *seip-1(tm4221)* mutants over two time periods from the beginning of adulthood. Each point represents a biological repeat. Statistical significances were determined by ordinary one-way ANOVA with *post hoc* Sidak’s test: ns: not significant. (E) Expression of wild-type SEIP-1 in the *seip-1(tm4221)* mutants decreases the embryonic lethality. Each point represents a biological repeat. Statistical significance was determined by two-tailed and unpaired t-test: \**P* < 0.05. (F) Confocal images of embryonic lipid droplets and yolk particles. Lipid droplets were stained with BODIPY and yolk particles were labelled by GFP-tagged lipoprotein VIT-2. Supersized lipid droplets in *seip-1(tm4221)* embryos are indicated by arrows. (G) Confocal images of wild-type and *seip-1(tm4221)* embryos with the plasma membrane labelled by GFP::PH(PLC1delta1) and the nucleus labelled by mCherry::Histone. Eggshells are indicated by dashed lines and gaps between eggshells and plasma membranes are indicated by asterisks.

Since the well-known function of seipin is to control lipid droplet homeostasis, we examined lipid droplets in embryos by BODIPY staining. Compared to wild-type embryos, there were several supersized lipid droplets in *seip-1(tm4221)* mutant embryos (Figure 1F). In contrast, there was no difference in yolk particles, labeled by the yolk protein VIT-2, between *seip-1(tm4221)* embryos and wild-type embryos. These results indicate that *seip-1(tm4221)* affects lipid droplet homeostasis.

To further characterize the phenotype of *seip-1(tm4221)* embryos, we labeled the plasma membrane and nucleus with fluorescent reporters. We found that *seip-1(tm4221)* embryos often contained multinucleate cells even at the early stage of embryogenesis (Figure 1G). To understand how these multinucleate cells formed, we used time-lapse microscopy to track cytokinesis. In *seip-1(tm4221)* embryos, cytokinesis proceeded more slowly and often aborted halfway through. Abrupt disruption of the adjoining plasma membrane, resulting in cell fusion, was also found in *seip-1(tm4221)* mutants (Figure S1A). In addition, the plasma membrane of cells in *seip-1(tm4221)* embryos was very close to the supporting outer eggshell, while there was a gap between the plasma membrane and the eggshell in wild-type embryos (Figure 1G). The eggshell is composed of six layers, among which the rigid chitin layer shapes the embryo and the permeability barrier layer maintains the internal osmotic pressure (Figure S1B) (Olson et al., 2012; Stein and Golden, 2018). The loss of space between the chitin layer and the plasma membrane in *seip-1(tm4221)* embryos indicates that the permeability barrier was disturbed, which caused hypo-osmotic swelling of the cells. Indeed, electron microscopy showed that the permeability barrier layer was missing in *seip-1(tm4221)* eggshells (Figure S1C). Accordingly, *seip-1(tm4221)* embryos swelled in hypotonic buffer and shrank in hypertonic buffer (Figure S1D). Moreover, the *seip-1(tm4221)* eggshell was permeable to DAPI dye (Figure S1E). These results suggest that *seip-1(tm4221)* embryos have an eggshell permeability defect, consistent with a previous report (Bai et al., 2020).

### *spin-4* and *nhr-114* suppress the embryonic lethality of *seip-1* mutants

To understand how *seip-1* affects eggshell integrity and embryogenesis, we carried out a genetic screen to identify suppressors of the highly penetrant embryonic lethal phenotype in *seip-1(tm4221)* mutants (Figure 2A). About 2,300 genomes were mutagenized by ethyl methanesulfonate (EMS). We identified two suppressors, *xd286* and *xd287*, which partially suppressed the embryonic lethality of *seip-1(tm4221)* mutants (Figure 2B and C). Single nucleotide polymorphism mapping and whole genome sequencing identified a missense mutation in *spin-4(xd286)* and a splicing site mutation in *nhr-114(xd287)* (Figure 2D and E). *spin-4* encodes a lysosomal/late-endosome transmembrane transporter of the major facilitator superfamily and its human ortholog SPNS1 may export sugar/sphingolipid (Rong et al., 2011). *nhr-114* encodes a transcription factor of the nuclear hormone receptor family and acts in the “B12-one-carbon cycle-PC (phosphatidylcholine)” axis (Giese et al., 2020).

**Fig. 2.**
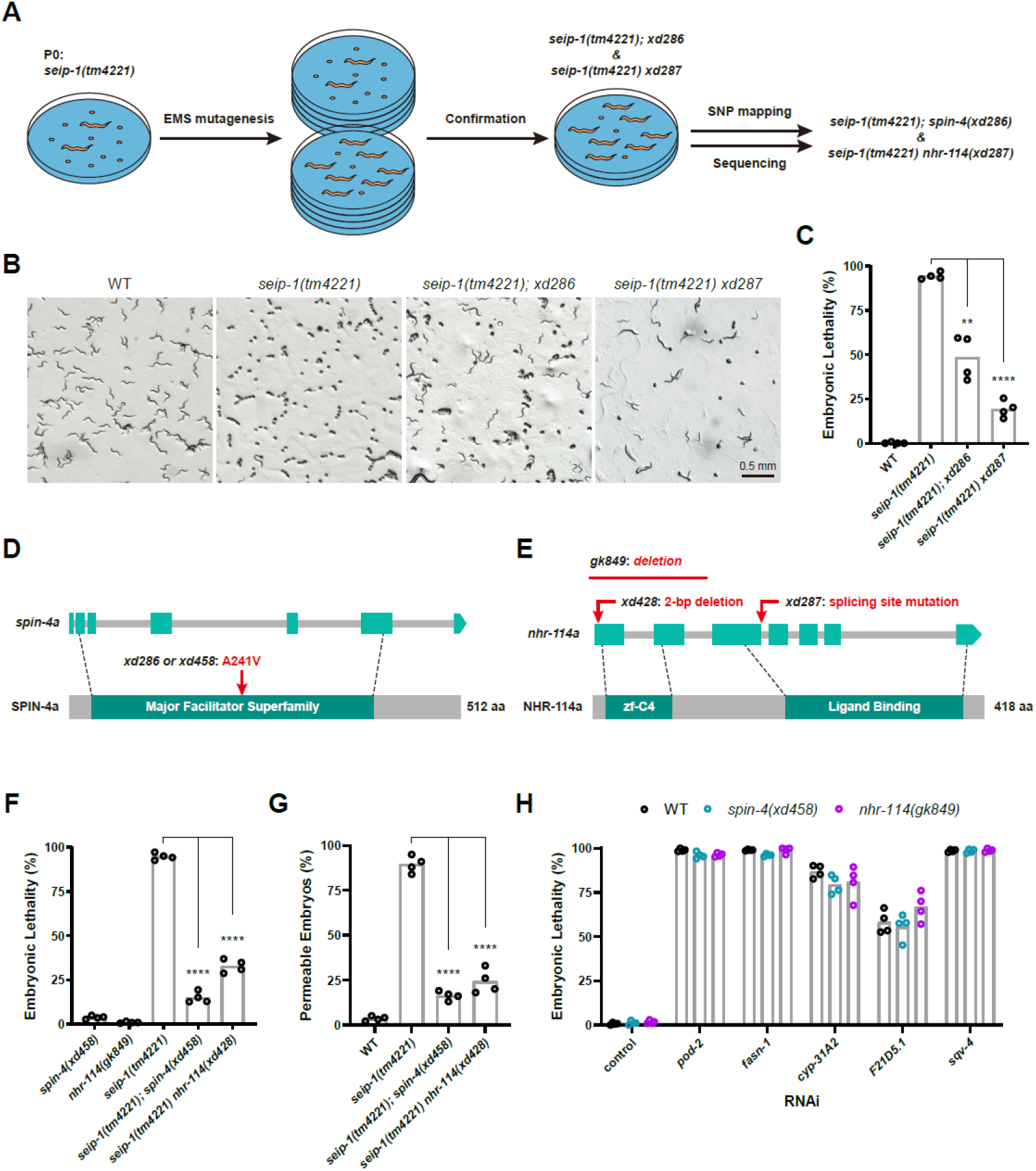
*spin-4* and *nhr-114* mutations suppress the embryonic lethality of *seip-1* mutants. (A) Workflow for the EMS mutagenesis screen and the identification of suppressor genes. The embryonic lethality phenotype was manifested by contrasting numbers of embryos and larvae on each plate. (B) Bright-field images of worm plates containing wild type, *seip-1(tm4221), seip-1(tm4221);xd286*, and *seip-1(tm4221) xd287* animals. (C) Quantification of the embryonic lethality of wild type, *seip-1(tm4221), seip-1(tm4221);xd286*, and *seip-1(tm4221) xd287.* Each point represents a biological repeat. Statistical significances were determined by Brown-Forsythe and Welch ANOVA with *post hoc* Dunnett’s T3 test: \*\**P* < 0.01, \*\*\*\**P* < 0.0001. (D) Diagram of the amino acid change in SPIN-4 isoform a caused by the *spin-4(xd286)* and *spin-4(xd458)* mutations. Light cyan rectangles indicate the exons and grey bars indicate the introns in the *spin-4* gene. The dark cyan rectangle highlights the conserved region of the major facilitator superfamily (MFS) in the SPIN-4 protein. Dashed lines indicate the start and end of the region encoding the conserved MFS domain. (E) Diagram of the molecular changes caused by the *nhr-114(xd287), nhr-114(xd428)* and *nhr-114(gk849)* mutations. Light cyan rectangles indicate the exons and grey lines indicate the introns in the *nhr-114* gene. Dark cyan rectangles highlight the DNA binding (zf-C4) and ligand binding domains in the NHR-114 protein. Dashed lines indicate the start and end of the regions encoding the two conserved domains. (F) Quantification of the embryonic lethality of mutant alleles. Each point represents a biological repeat. Statistical significances were determined by ordinary one-way ANOVA with *post hoc* Dunnett’s test: \*\*\*\**P* < 0.0001. (G) Quantification of the permeability of wild-type and mutant eggshells. The eggshell was judged to be permeable if the embryo shrunk in a hyperosmotic solution. Each point represents a biological repeat. Statistical significances were determined by ordinary one-way ANOVA with *post hoc* Dunnett’s test: \*\*\*\**P* < 0.0001. (H) Quantification of the embryonic lethality of wild type, *spin-4(xd458)* and *nhr-114(gk849)* animals fed on different RNAi bacteria.

To rule out the possibility that the suppression is caused by other background mutations, we generated *spin-4(xd458)* and *nhr-114(xd428)* mutations by CRISPR-Cas9. These mutations also partially suppressed the embryonic lethality of *seip-1(tm4221)* mutants (Figure 2F), which demonstrates that *spin-4* and *nhr-114* are *bona fide* genetic suppressors of *seip-1(tm4221)* embryonic lethality. Besides, *spin-4(xd458)* and *nhr-114(xd428)* also suppressed the permeable eggshell defect to a similar extent in *seip-1(tm4221)* embryos (Figure 2G), which indicates the importance of eggshell integrity to embryonic survival.

It was reported that seipin broadly affects cellular metabolism of lipids. Previous studies identified several genes involved in lipid or carbohydrate metabolism that are important for the formation of the eggshell, and deficiencies of these genes cause embryonic lethality similar to the *seip-1(tm4221)* mutation (Olson et al., 2012; Stein and Golden, 2018). Therefore, we tested the specificity of the suppression of *spin-4* and *nhr-114* mutations in *seip-1(tm4221)* mutants. Because *nhr-114* is closely linked to *seip-1*, we did not obtain a strain carrying the *nhr-114(xd428)* mutation alone. Instead, we used the deletion allele *nhr-114(gk849)* (Figure E). *spin-4(xd458)* and *nhr-114(gk849)* did not suppress the embryonic lethality induced by RNAi of genes involved in fatty acid synthesis (*pod-2* and *fasn-1*), fatty acid modification (*cyp-31A2*), or monosaccharide modification (*F21D5.1* and *sqv-4*) (Figure 2H). Put together, these results indicate that the *spin-4* and *nhr-114* mutations act as specific suppressors of the embryonic lethality of *seip-1(tm4221)* mutants.

### *spin-4* and *nhr-114* act in the same “B12-one-carbon cycle-PC” pathway

We then investigated how *spin-4* and *nhr-114* suppress the embryonic lethality of *seip-1(tm4221)* mutants. NHR-114 acts in the “B12-one-carbon cycle-PC” pathway (Figure 3A) (Giese et al., 2020). As a coenzyme, vitamin B12 regulates two important metabolic reactions in this axis (Figure 3A) (Giese et al., 2020). One reaction converts methylmalonyl-CoA to succinyl-CoA in the major propionate breakdown pathway. The second reaction, which is part of the one-carbon cycle, produces S-adenosylmethionine (SAM). SAM acts as a methyl donor to dozens of methyl receptors, including phosphoethanolamine, which is important for the synthesis of PC. When B12 is deficient, NHR-114 activates the expression of several genes in the B12 transport and one-carbon cycle pathway and promotes PC biosynthesis (Giese et al., 2020). *nhr-114* mutants displayed a diet-dependent sterility: they are sterile when growing on *E. coli* OP50, while they are fertile when growing on *E. coli* HT115 (Gracida and Eckmann, 2013). *E. coli* HT115 provides more B12 to *C. elegans* than *E. coli* OP50 (Watson et al., 2014), which explains the diet-dependent sterility phenotype of *nhr-114* mutants.

**Fig. 3.**
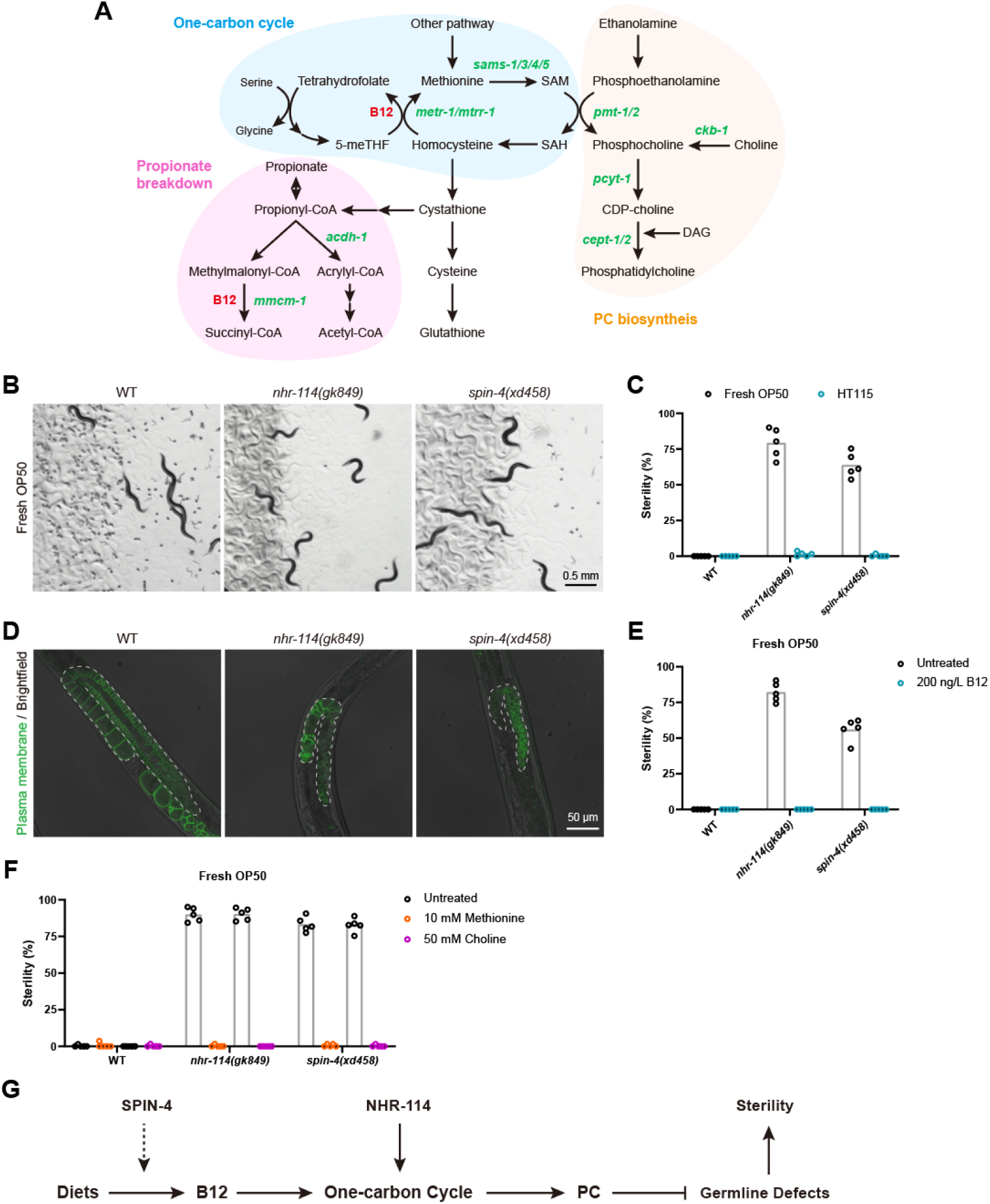
*spin-4* and *nhr-114* act in the “B12-one-carbon cycle-PC” axis. (A) Diagram of the association between the three metabolic pathways: propionate breakdown, one-carbon cycle and PC synthesis. CDP: cytidine 5’-diphosphocholine; 5-meTHF: 5-methyltetrahydrofolate; SAH: S-adenosylhomocysteine; SAM: S-adenosylmethionine; DAG: diacylglycerol. (B) Bright-field images of worm plates carrying wild type, *nhr-114(gk849)* mutants and *spin-4(xd458)* mutants on fresh OP50. Both mutants have a greatly decreased number of offspring (embryos and larvae). (C) Quantification of the sterility of wild type, *nhr-114(gk849)* mutants and *spin-4(xd458)* mutants grown on fresh OP50 or normal HT115 plates. (D) Confocal images of the germlines of day 1 adult wild type, *nhr-114(gk849)* mutants and *spin-4(xd458)* mutants labeled by GFP::PH(PLC1delta1). Dashed lines indicate the germline contours. (E and F) Quantification of the sterility of wild type, *nhr-114(gk849)* mutants and *spin-4(xd458)* mutants cultured on fresh OP50 supplied with or without 200 ng/L B12, 10 mM methionine or 50 mM choline. (G) Model of NHR-114 and SPIN-4 acting in the “B12-one-carbon cycle-PC” axis to regulate fertility.

The link between *spin-4* and *nhr-114* was previously unknown. We observed a similar semi-sterile phenotype when growing *nhr-114(gk849)* and *spin-4(xd458)* mutants on an OP50 diet (Figure S2A). Notably, the sterility is greatly increased when *nhr-114(gk849)* and *spin-4(xd458)* mutants were fed on a fresh OP50 diet (Figure 3B and C). Similar to *nhr-114(gk849), spin-4(xd458)* mutants are fully fertile when grown on an HT115 diet (Figure 3C). In addition, the germlines of *nhr-114(gk849)* and *spin-4(xd458)* mutants were distorted and smaller compared to wild type (Figure 3D). Taking the suppression of the *seip-1(tm4221)* mutation into account, these phenotypic similarities of *spin-4* and *nhr-114* mutants imply that the *spin-4* and *nhr-114* function in the same pathway.

We next explored whether *spin-4* functions in the “B12-one-carbon cycle-PC” axis. We examined the level of B12 with the widely used *Pacdh-1::GFP* reporter (Watson et al., 2014; Wei and Ruvkun, 2020). Expression of this reporter is inversely correlated with the organismal B12 level. Expression of *Pacdh-1::GFP* in wild-type animals was lower when the diet was normal HT115 compared to fresh OP50 (Figure S2B). Expression of *Pacdh-1::GFP* was increased in *spin-4(xd458)* mutants (Figure S2C and D). This result indicates that similar to *nhr-114, spin-4* mutants have a low organismal B12 level. Indeed, supplying B12 to fresh OP50 fully suppressed the sterility of *spin-4(xd458)* mutants (Figure 3E). Supplementation with methionine or choline also fully suppressed the sterility of *spin-4(xd458)* and *nhr-114(gk849)* mutants (Figure 3F and Figure S2E). These results demonstrate that *spin-4* functions in the “B12-one-carbon cycle-PC” axis (Figure 3G).

We next asked how *spin-4* regulates the “B12-one-carbon cycle-PC” axis. Extracellular B12 bound with a carrier protein is firstly taken up by the cell into its lysosome where the carrier protein is degraded; then free B12 is exported to cytosol by lysosomal ABC transporter for further modification before it functions as a coenzyme. It was reported that deficiency in lysosomal biogenesis or acidification causes B12 deficiency in *C. elegans* (Wei and Ruvkun, 2020). We speculated that SPIN-4, the *C. elegans* ortholog of the human lysosomal transporter SPNS1, may facilitate transport of B12 across the lysosomal. To examine the expression and protein localization of SPIN-4, we created a *Pspin-4::GFP* transcriptional fusion reporter and a *Pspin-4::spin-4::GFP* translational fusion reporter. The transcriptional GFP reporter was widely expressed, including in intestine and hypodermis (Figure S2F). The SPIN-4::GFP signal surrounded the signal from the lysosomal protease R07E3.1::mCherry reporter, which suggests that SPIN-4 is located on the lysosomal membrane (Figure S2G). Together, these results suggest that SPIN-4 likely facilitates lysosomal B12 transport and affects the “one-carbon cycle-PC” pathway.

### *nhr-114* and *spin-4* suppress the embryonic lethality of *seip-1(tm4221)* mutants through the “B12-one-carbon cycle-PC” axis

We then asked whether reducing the activity of the “B12-one-carbon cycle-PC” axis suppresses the embryonic lethality of *seip-1(tm4221)* mutants. Notably, fresh OP50 significantly decreased the embryonic lethal phenotype compared to normal OP50 and normal HT115 (Figure 4A). This result indicates that the embryonic lethality of *seip-1(tm4221)* mutants is diet-dependent. Since both SPIN-4 and NHR-114 regulate the “B12-one-carbon cycle-PC” axis, we supplied *seip-1(tm4221)* mutants with B12, methionine and choline. All of the metabolites fully blunted the alleviating effect of fresh OP50 on the embryonic lethality of *seip-1(tm4221)* mutants (Figure 4B). Importantly, supplementation of these metabolites also significantly reduced the suppressing effect of *nhr-114(xd428)* and *spin-4(xd458)* mutations on the embryonic lethality of *seip-1(tm4221)* mutants (Figure 4C). This suggests that the suppression effect of *nhr-114* and *spin-4* on *seip-1(tm4221)* mutants is indeed mediated through the “B12-one-carbon cycle-PC” axis.

**Fig. 4.**
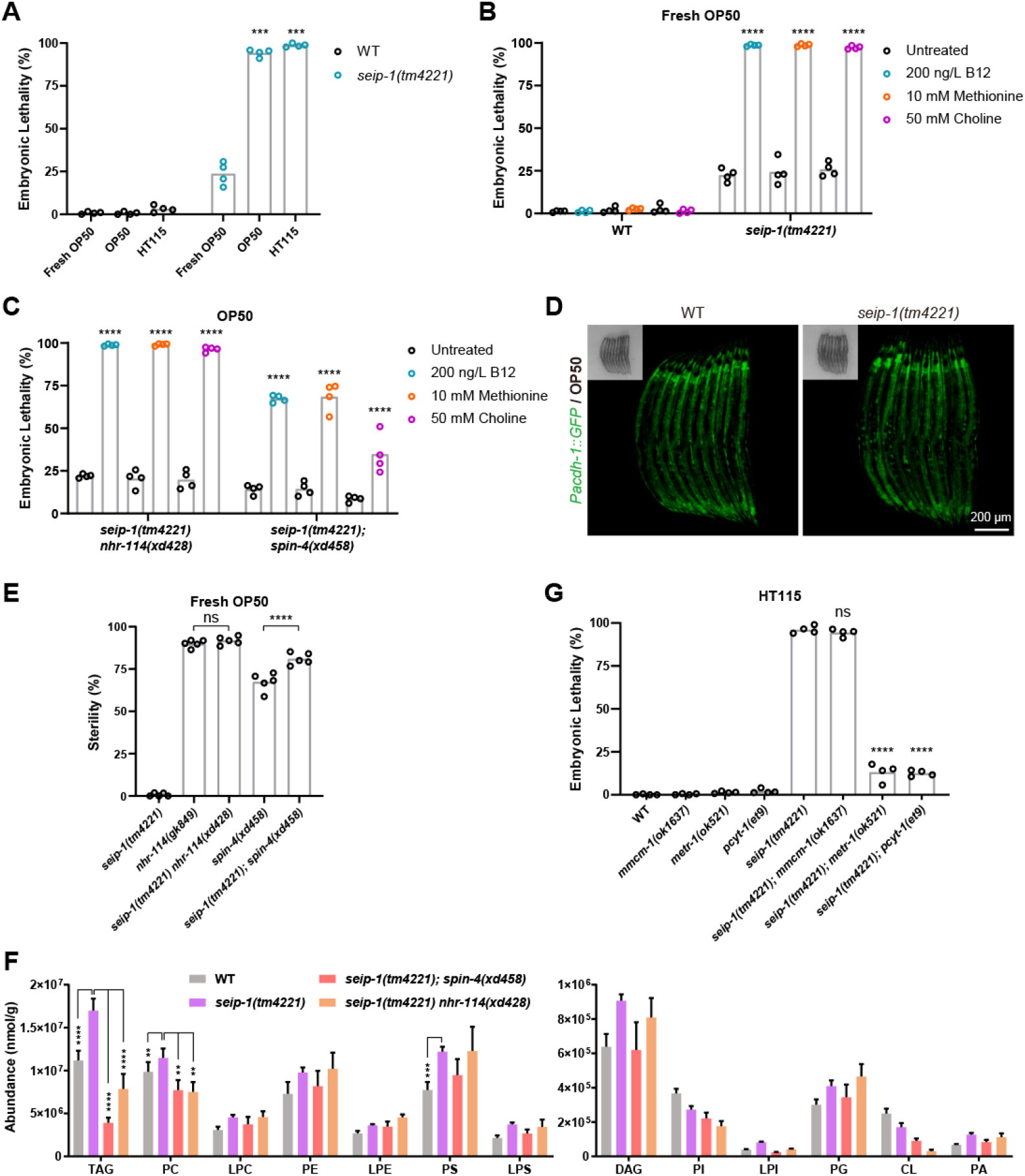
PC deficiency suppresses the embryonic lethality of *seip-1* mutants. (A) Quantification of the embryonic lethality of wild type and *seip-1(tm4221)* mutants fed on fresh OP50, normal OP50 or normal HT115. Each point represents a biological repeat. Statistically significant differences between *seip-1(tm4221)* mutants grown on fresh OP50 and each other diet were determined by Brown-Forsythe and Welch ANOVA with *post hoc* Dunnett’s T3 test: \*\*\**P* < 0.001. (B) Quantification of the embryonic lethality of wild type and *seip-1(tm4221)* mutants cultured on fresh OP50 supplied with or without 200 ng/L B12, 10 mM methionine or 50 mM choline. Each point represents a biological repeat. Statistically significant differences between *seip-1(tm4221)* mutants with and without each supplemented metabolite were determined by ordinary one-way ANOVA with *post hoc* Sidak’s test: \*\*\*\**P* < 0.0001. (C) Quantification of the embryonic lethality of *seip-1(tm4221) nhr-114(xd428)* and *seip-1(tm4221);spin-4(xd458)* double mutants cultured on fresh OP50 supplied with or without 200 ng/L B12, 10 mM methionine or 50 mM choline. Each point represents a biological repeat. Statistically significant differences between mutants with and without each supplemented metabolite were determined by two-way ANOVA with *post hoc* Tukey’s test: \*\*\*\**P* < 0.0001. (D) Confocal images of the *Pacdh-1::GFP* reporter in the wild-type or *seip-1(tm4221)* background. Insets are bright-field images. (E) Sterility was compared among different mutant animals fed on the fresh OP50 diet. Each point represents a biological repeat. Statistical significance was determined by ordinary one-way ANOVA with *post hoc* Sidak’s test: \*\*\*\**P* < 0.0001, ns: not significant. (F) Profiles of the lipid content in embryos from gravid adults of wild type, *seip-1(tm4221), seip-1(tm4221) nhr-114(xd428)* and *seip-1(tm4221);spin-4(xd458)*. Data was normalized to total protein. Error bars represent SEM. Statistically significant differences between *seip-1(tm4221)* and each other sample were determined by two-way ANOVA with *post hoc* Dunnett’s test: \*\**P* < 0.01, \*\*\**P* < 0.001, \*\*\*\**P* < 0.0001. (G) Quantification of the embryonic lethality of wild type and mutants fed on the normal HT115 diet. Each point represents a biological repeat. Statistically significant differences between the *seip-1(tm4221)* single mutant and each double mutant were determined by ordinary one-way ANOVA with *post hoc* Dunnett’s test: \*\*\*\**P* < 0.0001.

The suppression effect of PC prompted us to examine whether the activity of the “B12-one-carbon cycle-PC” axis is increased in *seip-1(tm4221)* mutants. We examined the expression of the *Pacdh-1::GFP* reporter in *seip-1(tm4221)* mutants, and found no difference compared to controls (Figure 4D). This suggests that *seip-1* deficiency does not affect the organismal B12 level. We also examined whether *seip-1(tm4221)* mutation affects the sterility of *nhr-114* and *spin-4* mutants. The sterility of *nhr-114(xd428)* and *spin-4(xd458)* mutants was not suppressed by the *seip-1(tm4221)* mutation (Figure 4E). Therefore, the activity of the “B12-one-carbon cycle-PC” axis is probably not increased in *seip-1(tm4221)* mutants.

To examine the changes in PC levels associated with suppression of the *seip-1(tm4221)* mutation, we performed lipid profiling to measure the levels of PC and other lipids in *seip-1(tm4221)* single mutant embryos and in *seip-1(tm4221) nhr-114(xd428)* and *seip-1(tm4221);spin-4(xd458)* double mutant embryos. Compared to wild type, the levels of diacylglycerol (DAG) and triacylglycerol (TAG) were dramatically increased in *seip-1(tm4221)* mutant embryos (Figure 4F). The most abundant phospholipids, PC, PE and PS, were also slightly increased in *seip-1(tm4221)* mutants. In *seip-1(tm4221) nhr-114(xd428)* and *seip-1(tm4221);spin-4(xd458)* double mutants, the levels of PE, PS and other lipids were not changed, compared to *seip-1(tm4221)* alone (Figure 4F). Consistent with the notion that suppression of the *seip-1(tm4221)* mutation occurs by reducing the activity of the “B12-one-carbon cycle-PC” axis, the level of PC was significantly decreased in both *seip-1(tm4221) nhr-114(xd428)* and *seip-1(tm4221);spin-4(xd458)* double mutants (Figure 4F). In addition, the level of TAG was also decreased in these double mutants, compared to *seip-1(tm4221)* alone. In line, the mutation *metr-1(ok521)*, which affects the one-carbon cycle, and the mutation *pcyt-1(et9)*, which affects PC synthesis, significantly suppressed the embryonic lethality of *seip-1(tm4221)* mutants, while the mutation *mmcm-1(ok1637)*, which affects the propionate breakdown pathway, did not (Figure 4G). Put together, these results support the idea that lowering the activity of the “B12-one-carbon cycle-PC” axis suppresses the embryonic lethality of *seip-1(tm4221)* mutants.

### Suppression of the embryonic lethality of *seip-1* mutants by PC deficiency depends on PUFAs

We next explored the underlying mechanism of the suppression of PC deficiency on *seip-1(tm4221)* embryonic lethality. Recent studies reported that dietary supplementation of GLA(C18:3n-6) and DGLA(C20:3n-6), two ω-6 polyunsaturated fatty acids (PUFAs), promote the enrichment of SEIP-1 in an ER subdomain and partially rescues the embryonic lethality of *seip-1* mutants (Bai et al., 2020; Cao et al., 2019). To investigate whether *nhr-114* and *spin-4* mutations suppressed the embryonic lethality of *seip-1(tm4221)* mutants by increasing the level of PUFAs, we analyzed the abundance of free fatty acids (FFAs). The levels of FFAs were increased in *seip-1(tm4221)* embryos and decreased in *seip-1(tm4221) nhr-114(xd428)* and *seip-1(tm4221);spin-4(xd458)* double mutant embryos (Figure 5A). Among FFAs, we also compared the relative levels of saturated fatty acids (SFAs), monounsaturated fatty acids (MUFAs) and polyunsaturated fatty acids (PUFAs). While SFA and MUFA levels were not changed in *seip-1(tm4221)* mutants compared to wild type, PUFA levels were increased in *seip-1(tm4221)* embryos. In *seip-1(tm4221) nhr-114(xd428)* and *seip-1(tm4221);spin-4(xd458)* double mutant embryos, SFA levels were increased and MUFA and PUFA levels were decreased compared to *seip-1(tm4221)* alone (Figure 5B). The abundance of C20:3 fatty acids were dramatically increased in *seip-1(tm4221)* mutants. In both double mutant embryos, the abundance of C20:3 fatty acids were dramatically decreased compared to *seip-1(tm4221)* single mutants (Figure 5A). In addition, the abundance of most FFAs was unchanged or even decreased in *nhr-114(gk849)* and *spin-4(xd458)* single mutants (Figure 5C). These data suggest that the suppression effect of *nhr-114* and *spin-4* mutations on the embryonic lethality of *seip-1(tm4221)* is unlikely to occur through increasing the levels of DGLA or other PUFAs.

**Fig. 5.**
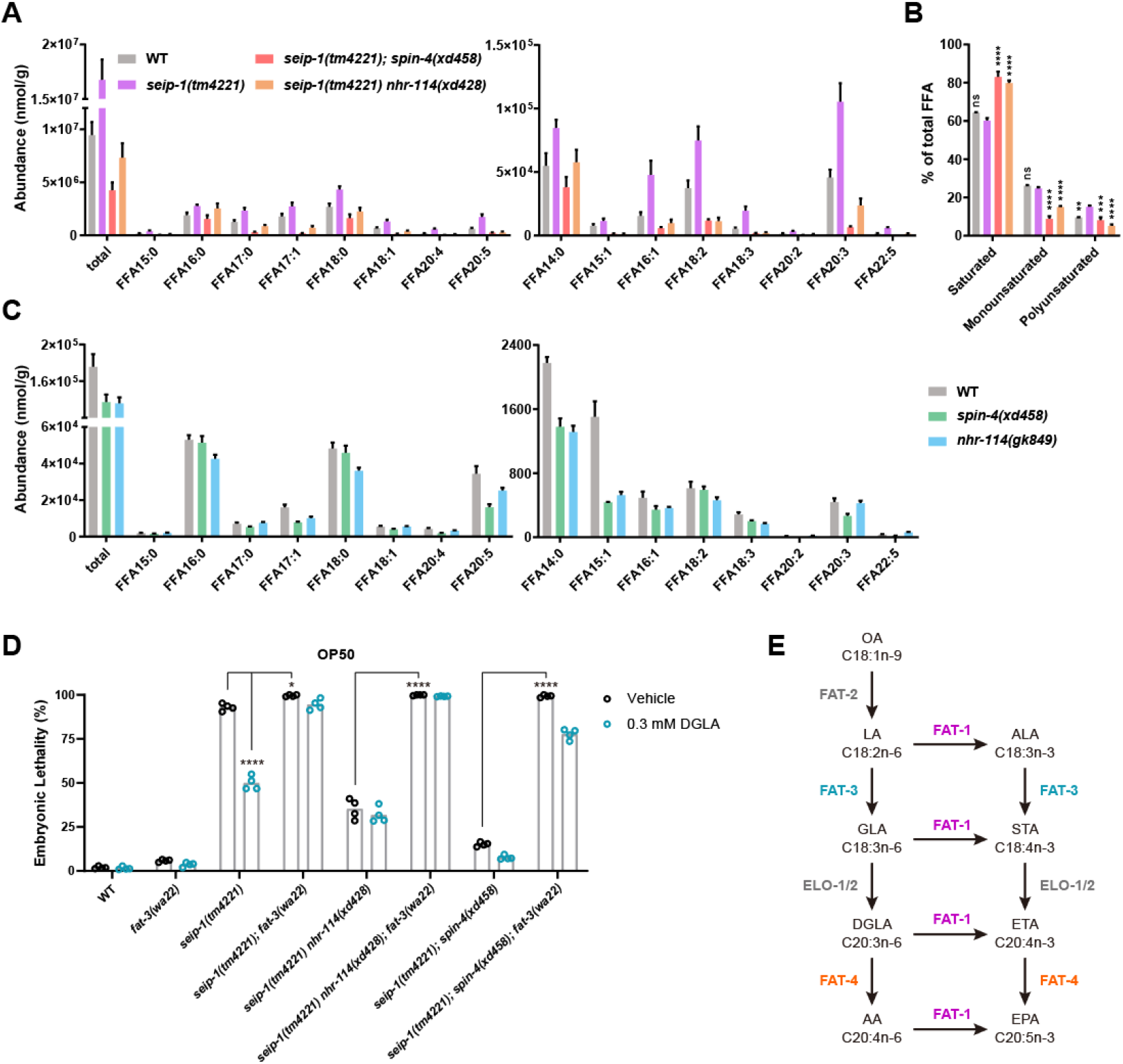
Suppression of the embryonic lethality of *seip-1* mutants by PC deficiency depends on PUFAs. (A) Profiles of free fatty acids (FFAs) in early embryos of wild type, *seip-1(tm4221), seip-1(tm4221) nhr-114(xd428)* and *seip-1(tm4221);spin-4(xd458)*. Data was normalized to total protein. Error bars represent SEM. (B) Relative levels to total FFAs of saturated FFAs, monounsaturated FFAs and polyunsaturated FFAs in embryos from gravid adults of wild type, *seip-1(tm4221), seip-1(tm4221) nhr-114(xd428)* and *seip-1(tm4221);spin-4(xd458)*. Error bars represent SEM. Statistically significant differences between *seip-1(tm4221)* and each other sample were determined by two-way ANOVA with *post hoc* Dunnett’s test: \*\**P* < 0.01, \*\*\**P* < 0.001, \*\*\*\**P* < 0.0001. (C) Profiles of FFAs in day 1 adults of wild type, *spin-4(xd458)* mutants and *nhr-114(gk849)* mutants. Data was normalized to total protein. Error bars represent SEM. (D) Quantification of the embryonic lethality of wild type and mutants fed on the OP50 diet supplied with 0.3 mM DGLA or vehicle. Each point represents a biological repeat. Statistical significances were determined by two-way ANOVA with *host hoc* Tukey’s test: \**P* < 0.05, \*\*\*\**P* < 0.0001. (E) The pathway for fatty acid elongation and desaturation in *C. elegans*. Enzymes that catalyze each step are indicated alongside the arrows. POA: palmitoleic acid; VA: vaccenic acid; PA: palmitic acid; SA: stearic acid; OA: oleic acid; LA: linoleic acid; ALA: alpha-linolenic acid; GLA: gamma-linoleic acid; STA: stearidonic acid; DGLA: dihomogamma-linolenic acid; ETA: eicosatetraenoic acid; AA: arachidonic acid; EPA: eicosapentaenoic acid.

We further examined the relationship between *nhr-114/spin-4*-mediated suppression and PUFAs. Consistent with a previous report (Bai et al., 2020), DGLA supplementation partially suppressed the embryonic lethality of *seip-1(tm4221)* mutants (Figure 5D). Interestingly, DGLA supplementation did not enhance the suppression effect of the *nhr-114(xd428)* on *seip-1(tm4221)* mutations. The suppression effect of the *spin-4(xd458)* mutation on *seip-1(tm4221)* mutants was slightly enhanced by DGLA supplementation (Figure 5D). *De novo* PUFA biosynthesis starts from the desaturation of oleic acid by FAT-2, followed by three other desaturases, FAT-1, FAT-3 and FAT-4, to generate complex PUFAs (Kahn-Kirby et al., 2004) (Figure 5E). The *fat-3(wa22)* mutation, which blocks the synthesis of downstream PUFAs including DGLA, enhanced the embryonic lethality of *seip-1(tm4221)* mutants and blocked the rescue effect of DGLA supplementation (Figure 5D). Notably, the *fat-3(wa22)* mutation completely abolished the suppression effect of *nhr-114* and *spin-4* mutations on the embryonic lethality of *seip-1(tm4221)* mutants (Figure 5D). Together, these results indicate that the rescuing effect of PC deficiency on the embryonic lethality of *seip-1(tm4221)* mutants depends on PUFAs.

### PC deficiency enhances the lipid droplet phenotype of *seip-1* mutants

The suppression effect of *spin-4(xd458)* and *nhr-114(xd428)* mutations on the embryonic lethality of *seip-1(tm4221)* mutants provides a great opportunity to address a longstanding question about seipin: what is the relationship between its roles in cellular lipid droplet homeostasis and physiological function (i.e., embryogenesis in this study). In mature oocytes (Figure 6A), *seip-1(tm4221)* mutants exhibit numerous large lipid droplets compared to wild type (Figure 6B and C). This “supersized” lipid droplet phenotype is also found in yeast, fly and mouse seipin mutants (Cui et al., 2011; Fei et al., 2008; Fei et al., 2011b; Pagac et al., 2016; Szymanski et al., 2007; Tian et al., 2011).

**Fig. 6.**
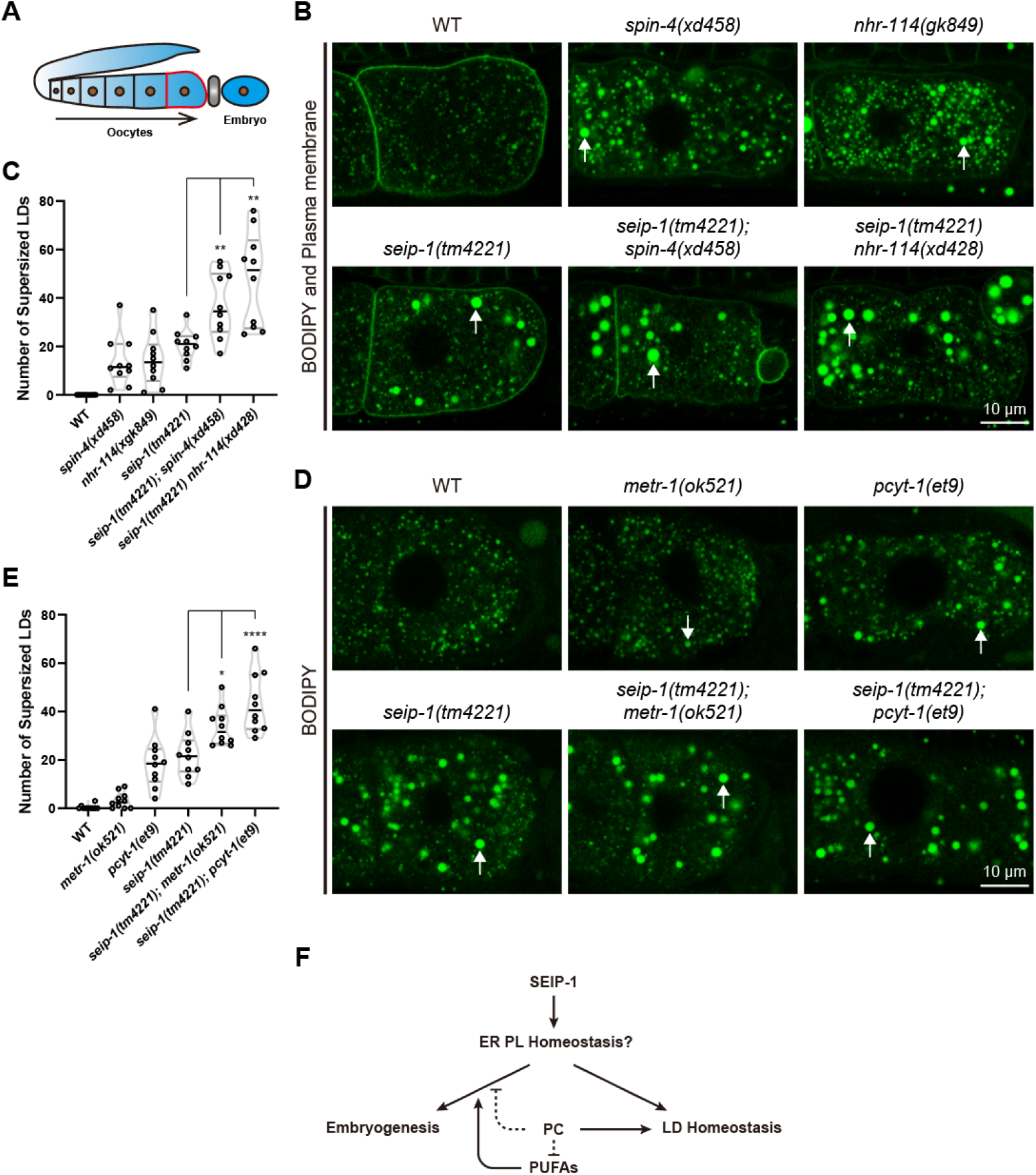
PC deficiency enhances the lipid droplet phenotype of *seip-1* mutants. (A) Diagram of the development of *C. elegans* oocytes. The plasma membrane of a mature oocyte that is ready for fertilization is displayed as red. The spermatheca is indicated in grey. (B) Representative confocal images of supersized lipid droplets in mature oocytes of wild type and mutants. White arrows indicate supersized lipid droplets. (C) Quantification of the supersized lipid droplets in B. Each point represents a biological repeat. Statistically significant differences between the *seip-1(tm4221)* single mutant and each double mutant were determined by Brown-Forsythe and Welch ANOVA with *post hoc* Dunnett’s T3 test: \*\**P* < 0.01. (D) Representative confocal images of supersized lipid droplets in mature oocytes of wild type and mutants. White arrows indicate supersized lipid droplets. (E) Quantification of the supersized lipid droplets in D. Each point represents a biological repeat. Statistically significant differences between the *seip-1(tm4221)* single mutant and each double mutant were determined by ordinary one-way ANOVA with *post hoc* Dunnett’s test: \**P* < 0.05, \*\*\*\**P* < 0.0001. (F) Working model for the role of PC, PUFAs and SEIP-1 in the regulation of embryogenesis and lipid droplet homeostasis. ER: endoplasmic reticulum; PL: phospholipid; PC: phosphatidylcholine; PUFA: polyunsaturated fatty acid; LD: lipid droplet.

We then examined the effect of *spin-4(xd458)* and *nhr-114(xd428)* mutations on lipid droplets in *seip-1(tm4221)* mutant oocytes. Similar to *seip-1(tm4221)* mutants, *spin-4(xd458)* and *nhr-114(xd428)* mutants exhibit large lipid droplets (Figure 6B and C), consistent with previous findings that PC deficiency leads to large lipid droplets (Krahmer et al., 2011; Walker et al., 2011; Wang et al., 2018). Remarkably, *spin-4(xd458)* and *nhr-114(xd428)* mutations further increased the number of large lipid droplets in *seip-1(tm4221)* oocytes (Figure 6B and C). In line with that, the mutants *metr-1(ok521)* and *pcyt-1(et9)*, which are defective in PC synthesis, also display large lipid droplets. The number of large lipid droplets is also significantly increased in *seip-1(tm4221);metr-1(ok521)* and *seip-1(tm4221);pcyt-1(et9)* double mutants compared to *seip-1(tm4221)* single mutants (Figure 6D and E). Therefore, while PC deficiency suppresses the embryonic lethality in *seip-1(tm4221)* mutants, it exacerbates the large lipid droplet phenotype in *seip-1(tm4221)* mutants. These results suggest that seipin may regulate embryogenesis and lipid droplet biogenesis through distinct mechanisms (Figure 6F).

## Discussion

In *seipin-deficient* organisms, the causal link between the cellular defect in lipid droplet homeostasis and the physiological defects is not clear. This study reveals that *seip-1* embryonic lethality is suppressed by reducing PC synthesis. *nhr-114*, a known regulator of the “B12-one carbon cycle-PC” axis, and *spin-4*, a new player in that axis, were identified as *seip-1* suppressors. The choline-phosphate cytidylyltransferase mutant *pcyt-1*, which is defective in PC synthesis, also suppresses the embryonic lethality of *seip-1* mutants. Consistent with previous reports (Fei et al., 2011b; Krahmer et al., 2011; Pagac et al., 2016), both *seip-1* mutation and PC deficiency mutations result in large lipid droplets. Interestingly, while PC deficiency suppresses *seip-1* embryonic lethality, it enhances the large lipid droplet phenotype of *seip-1* mutants. Therefore, our results suggest seipin-mediated embryogenesis is independent of lipid droplet homeostasis (Figure 6F).

The suppression of *seip-1* embryonic lethality by reduction of PC is unexpected. PC is a major phospholipid on lipid droplets. While reducing PC level results in large lipid droplets from lipid droplet coalescence (Krahmer et al., 2011), the formation of lipid droplets is also an adaptation to PC deficiency (Koh et al., 2018; Vevea et al., 2015). *Seipin* mutation also results in aberrant lipid droplet biogenesis with the formation of large lipid droplets, a phenotype similar to PC deficiency. Therefore, it is quite surprising that reducing PC synthesis suppresses the physiological defect of *seip-1* mutants. Similar to PC, the link between PUFA and lipid droplet dynamics is also obvious (Romanauska and Köhler, 2021; Xie and Roy, 2015). Notably, PUFA affects lipid droplet diversity by controlling the localization of seipin (Cao et al., 2019). Although the PUFA composition is changed in mouse and worm *seipin* mutants (Bai et al., 2020; Cao et al., 2019; Chen et al., 2013), this cannot explain the suppression of the *seip-1* mutant phenotype by PUFA supplementation. Besides, inhibiting PUFA synthesis represses LD defects induced by hepatic seipin deficiency (Lounis et al., 2017). Together, PUFA is likely to have opposite actions on the molecular and physiological defects induced by seipin deficiency. The suppression of *seip-1* embryonic lethality by PC reduction requires PUFA. The mechanistic link between PC and PUFA is not clear here. It is possible that PC and PUFA act in parallel, but converge on the same target/process. Alternatively, PC may directly inhibit PUFA function (Figure 6F).

We propose two possibilities to reconcile the lipid droplet homeostasis function and embryogenesis function of seipin. First, seipin may be multifunctional. Besides its role in lipid droplet biogenesis, seipin may function in other processes. In line with that, yeast seipin inhibits sphingolipid biogenesis. This function is parallel to its role in lipid droplet biogenesis, because inhibition of sphingolipid biogenesis only has a minor effect on lipid droplet biogenesis (Su et al., 2019). The diverse functions of seipin may be mediated by its binding with different partners (Ding et al., 2018; Pagac et al., 2016; Su et al., 2019; Yang et al., 2014). Another possibility is that seipin acts in a step before lipid droplet homeostasis and this step is shared with both lipid droplet homeostasis and embryogenesis/lipid barrier synthesis (Figure 6F). Consistent with this hypothesis, biogenesis of lipid droplets that contain only retinyl or steryl esters does not depend on seipin (Molenaar et al., 2021), which suggests that seipin does not directly determine droplet formation. The shared function could be in ER phospholipid homeostasis (Fei et al., 2011a; Gao et al., 2019).

It is possible that seipin regulates ER phospholipid homeostasis to counteract local phospholipid imbalances, membrane curvature changes or thermodynamic instability of neutral lipids in lipid bilayers to facilitate lipid droplet formation and lipid barrier synthesis/formation in an ordered and regulated way. Seipin may affect ER membrane lipid homeostasis by acting as a phospholipid synthetase/lipid transporter/flippase/scramblase or as a modulator of these functions (Gao et al., 2019; Ghanbarpour et al., 2021; Li et al., 2021; Zhao et al., 2017). For example, Seipin binds to GPAT3/4 (glycerol-3-phosphate acyltransferase) and negatively modulates its activity. Seipin deficiency leads to increased levels of phosphatidic acid, a conical membrane phospholipid, which promotes the negative curvature of membranes (Fei et al., 2011b; Pagac et al., 2016; Tian et al., 2011). Our previous study identified seipin as a partner of SERCA (sarcoplasmic-endoplasmic reticulum calcium ATPase) (Bi et al., 2014). Interestingly, VMP1, a phospholipid scramblase, is also a partner of SERCA (Li et al., 2021; Zhao et al., 2017).

The suppression of the embryonic lethality is likely due to recovery of the lipid barrier in the eggshell (Olson et al., 2012; Stein and Golden, 2018). It is possible that seipin-regulated ER membrane lipid balance is required for lipid barrier construction (the synthesis of specific lipids or the formation of the barrier). The altered ER lipid balance caused by *seipin* deficiency can be partially compensated by PC reduction or PUFA supplementation. PC is a membrane-forming cylindrical phospholipid, and therefore reducing PC affects membrane packing, permeability, and intrinsic membrane curvature as well as inducing acyl chain remodeling of the remaining phospholipids (Boumann et al., 2006; Haider et al., 2018; Li et al., 2006; Naito et al., 2015; Vanni et al., 2014). Long chain PUFAs are conformationally flexible and their incorporation into phospholipids greatly changes the physicochemical properties of membranes. PC reduction and PUFA supplementation may remodel the membrane lipid composition and change membrane properties to facilitate eggshell lipid barrier construction.

There have been several attempts to restore the normal physiological behaviors of *seipin*-deficient mice. These studies either directly target different physiological defects associated with *seipin* deficiency or originate from suppression of the lipid droplet phenotype (Dollet et al., 2016; Gao et al., 2020; Xu et al., 2015). Our finding raises the possibility that the well-studied cellular lipid droplet defect and physiological defects caused by *seipin* deficiency may be separable. The autonomous action of seipin supports this notion (McIlroy et al., 2020; McIlroy et al., 2018). An immediate application is to treat *seipin* deficiency with an inhibitor of PC synthesis. In addition, dietary manipulation of the B12, methionine or choline level is another appealing intervention for physiological defects in BSCL2 patients. The results may be beneficial for future treatment of diseases associated with *seipin* mutations and may extend to other diseases.

## Material and methods

### Strains and maintenance

The *C. elegans* strains used in this study are listed in table S1. Bristol N2 was used as wild-type control. All *C. elegans* strains were cultured at 22 °C on nematode growth medium (NGM) plates. *E. coli* OP50 was seeded onto fresh NGM plates for 2 days to allow lawn formation; this was defined as the “fresh OP50” diet. *E. coli* OP50 and *E. coli* HT115 were seeded onto fresh NGM plates for 7 days; these were defined as the “normal OP50 or HT115” diets. The diets can be stored at 4 °C for no more than 1 week before utilization.

### EMS screen and suppressor identification

For the suppressor screen, P0 L4 larvae of *seip-1(tm4221)/nT1[qIs51]* mutants were mutagenized by EMS. Every 3 F1 larvae were transferred to a new OP50 plate, and every 10 F2 larvae that no longer contained balancer fluorescence were cultured on a new OP50 plate. F2 plates that contained obviously increased numbers of F3 larvae were kept. The putative suppressors from F2 plates were transferred to new OP50 plates and were selected for increased progeny numbers through several generations.

To map the chromosomal location of the suppressors, the suppressors were crossed with the CB4856 strain and each F2 animal was cultured alone on an OP50 plate. F2 animals that were homozygous for the *seip-1(tm4221)* mutation were classified into two group according to their progeny number. Then single nucleotide polymorphism mapping was applied to these two groups as described (Davis et al., 2005).

The genomic DNA of suppressor-containing strains was extracted for whole-genome sequencing. The candidate suppressor genes were selected according to the mapping and sequencing results. To knock out *nhr-114* via CRISPR-Cas9 technology, two sgRNA sequences targeting to the first exon of *nhr-114* were designed and validated for specificity through BLAST searching against the whole genome. The sgRNA sequences were inserted into pDD162 plasmid and sequenced for validity. Due to the close linkage of *nhr-114* and *seip-1*, the recombinant plasmids were microinjected into *seip-1(tm4221)* mutant animals. Offspring were singled to examine the mutation status by sequencing. The homozygous mutant, *seip-1(tm4221) nhr-114(xd428)*, was obtained in next generation along with the loss of the Cas9 plasmid. A point mutation in *spin-4(xd458)* was generated from wild-type animals through CRISPR-Cas9 by SunyBiotech company. Then the *spin-4(xd458)* mutation was crossed into *seip-1(tm4221)* mutant to generate *seip-1(tm4221);spin-4(xd458)* double mutants.

### Molecular biology

The genomic DNA of *seip-1* was amplified from N2 genomic DNA. The forward primer was 5’-catcacgtgttcgctcgctgg-3’ and the reverse primer was 5’-tgccgacgaggacggttcgac-3’. The transgenic worm *seip-1(tm4221);xdEx1640(Pseip-1::seip-1 + rol-6[su1006])* was generated by microinjecting the amplified *seip-1* genomic DNA with a co-injection marker. Approximately 3.5 kb of upstream sequence of *spin-4* was cloned from wild-type genomic DNA and inserted into the multiple cloning site of pSM plasmid to generate the *spin-4* promoter fusion GFP reporter *Pspin-4::GFP.* The reporter was expressed in wild-type animals by microinjection with a co-injection marker to generate *xdEx2409 [Pspin-4::GFP + Podr-1::RFP]* animals. The coding sequence of SPIN-4 isoform a was cloned from wild-type cDNA and inserted in-frame between the *vha-6* promoter and GFP-coding sequence of *pSM::Pvha-6::GFP* through *in vitro* recombination to generate the *Pvha-6::spin-4::GFP* reporter. The reporter was expressed in *qxIs448 [R07E3.1::mCherry]* animals that express a mCherry-labelled lysosomal peptidase by microinjection with a co-injection marker to generate *qxIs448 [R07E3.1::mCherry];xdEx2483 [Pvha-6::SPIN-4::GFP + Podr-1::RFP]* animals.

### Quantification of physiological phenotypes

To quantify embryonic lethality, synchronized L1 larvae were cultured to day 1 adults (24 hours post mid-L4 stage). For each genotype or treatment, four independent plates were cultured. 20-30 adult animals per plate were picked to a new plate and allowed to lay embryos for 2-3 hours. Then the animals were removed and the total number of embryos was counted. After 20 hours, the dead embryos were counted. Embryonic lethality was calculated as the percentage of dead embryos among total embryos. To determine brood size, a single L4 animal was transferred to a new OP50 plate. Every 24 hours since the start of adulthood, the animal was transferred to a new plate and the embryos and larvae on the previous plate were counted. For each genotype, 10-20 L4 animals were singled for the analysis. To quantify sterility, synchronized L1 larvae were cultured to day 1 adults. For each genotype or treatment, five independent plates were cultured. More than 50 adults were picked per plate for inspection of *in utero* embryos under differential interference contrast microscopy (DIC). Sterile animals were characterized by an empty uterus. Sterility was calculated as the percentage of sterile animals among total animals.

Hypertonic solution (250 mM KCl, 5 mM HEPES) or hypotonic solution (100 mM KCl, 5 mM HEPES) was used to identify the integrity of the eggshell. Embryos were released from gravid adults by cutting the animals with two 1 mL syringe needles in the tested solution within an artificial hole in a 2.5% agarose pad to relieve the pressure from coverslip. Representative DIC images of 2-cell embryos were captured by ZEISS microscopy. Eggshell permeability was quantified by calculating the percentage of shrunken embryos among total embryos in hypertonic solution. For each genotype, 4 repeats were performed, each with 100 embryos in total.

### Staining and imaging

For DAPI staining, embryos were directly released from gravid adults in a droplet of DAPI (ThermoFisher Scientific) solution (2 μg/mL in M9 buffer) on a microscope slide. The slides were kept in wet box and stained for 20 minutes in the dark. To visualize lipid droplets in embryos, the embryos released from gravid adults were stained in BODIPY 493/503 (ThermoFisher Scientific) solution (5 μg/mL in M9 buffer) for 20 minutes. After one wash with M9 buffer, stained embryos were transferred onto a 2.5% agarose pad and sealed by a coverslip. The samples were immediately imaged by fluorescence microscopy or confocal microscopy. To visualize yolk particles in embryos, the embryos were immediately imaged by confocal microscopy after being released from gravid adults.

To visualize lipid droplets in oocytes, synchronized day 1 adults were washed off from culture plates with M9 buffer. The supernatant was discarded, then the animals were resuspended in BODIPY solution and stained in the dark for 3 hours with gentle shaking. The animals were then transferred to a new food plate to recover for 3 hours. The lipid droplets in mature oocytes were imaged by confocal microscopy. The numbers of supersized lipid droplets (diameter > 1 μm) were counted over a 14-μm z-axis thickness per mature oocyte. Ten oocytes were analyzed for each genotype.

To image embryos labelled by reporters for the plasma membrane and nucleus, embryos were released from gravid adult animals in a drop of M9 buffer. The embryos were transferred onto a 2.5% agarose pad and a coverslip was lightly placed over them. Next, the edge of the coverslip was sealed by wax and the gaps around the agarose pad were filled with egg salt buffer. Then the embryos were imaged by confocal microscopy (Leica SP8).

To visualize the expression of genes in adults, synchronized L1 larvae expressing the *Pacdh-1::GFP* reporter were cultured to day 1 adults. For each genotype, 10 adult animals were picked out for confocal microscopy at a 2.5 μm z-axis interval. The final images were created by merging all the z-axis photos covering the whole animals. To quantify the expression of the *Pacdh-1::GFP* reporter in wild type and *spin-4(xd458)* mutants, three parts of the images was segmented to measure the mean intensity by ImageJ software.

To analyze the subcellular location of SPIN-4, the intestine of adult *qxIs448 [R07E3.1::mCherry];xdEx2483 [Pvha-6::SPIN-4::GFP + Podr-1::RFP]* animals was imaged by confocal microscopy. To highlight the relative location of SPIN-4 and R07E3.1, line profiles of the intensity of both GFP and mCherry were analyzed by ImageJ software.

### Electron microscopy

Day 1 adult animals were collected for high-pressure freezing EM imaging as described (Yang et al., 2020). 60-nm ultrathin sections were prepared and imaged at 80 kV using a JEM-1400 TEM (Hitachi HT7700) with a Gatan832 4k × 2.7k CCD camera.

### RNAi feeding treatment

The RNAi feeding treatment was carried out as described (Liu et al., 2014). The synchronized L1 larvae were seeded onto RNAi plates and cultured to day 1 adults to determine the embryonic lethality.

### Metabolite supplementation

B12 (Sigma-Aldrich) was dissolved and diluted in ddH_2_O to make a 200 μg/L stock solution. NGM medium containing 200 ng/L B12 was made by adding 1 mL stock solution into 1 L sterile NGM before pouring plates. NGM medium containing 10 mM methionine or 50 mM choline was made by directly adding 1.5 g L-methionine (BioDee Biotechnology) or 7 g choline chloride (Sigma-Aldrich) respectively into 1 L sterile NGM before pouring plates. NGM medium containing 300 μM DGLA (Cayman Chemicals) or 1:1000 ethanol (as control) was made by adding 1 mL DGLA stock (300 mM in ethanol) or 1 mL ethanol respectively into 1 L sterile NGM before pouring plates. NGM plates containing DGLA were stored in the dark.

### Lipid profiling analysis

Animal collection: About 20,000 synchronized day 1 adults were washed off from culture plates by M9 buffer. The animals were suspended in M9 buffer to digest the food for 1 hour and then washed another 3 times with M9 buffer. The animal pellets were stored at −80 °C until lipid analysis. Embryo collection: About 50,000 synchronized day 1 adults were washed off from culture plates by M9 buffer. The animals were lysed by bleach buffer (1.25 M NaOH, 25% v/v bleach) to release the embryos. Then the embryo pellets were washed 3 times before being stored at −80 °C until lipid analysis.

Lipids were extracted using a modified version of the Bligh and Dyer’s method as described previously (Lam et al., 2017). Briefly, tissues were homogenized in 750 μL of chloroform:methanol 1:2 (v/v) with 10% deionized water on a Bead Ruptor (Omni, USA). The homogenate was then incubated at 1500 rpm for 1 h at 4°C. At the end of the incubation, 350 μL of deionized water and 250 μL of chloroform were added to induce phase separation. The samples were then centrifuged and the lower organic phase containing lipids was extracted into a clean tube. Lipid extraction was repeated once by adding 500 μL of chloroform to the remaining tissues in aqueous phase, and the lipid extracts were pooled into a single tube and dried in the SpeedVac under OH mode. Samples were stored at −80 □ until further analysis.

Polar lipids were analyzed using an Agilent 1260 HPLC system coupled with a triple quadrupole/ion trap mass spectrometer (5500 Qtrap; SCIEX) (Lam et al., 2021). Separation of individual lipid classes of polar lipids by normal phase (NP)-HPLC was carried out using a Phenomenex Luna 3 μm-silica column (internal diameter 150 × 2.0 mm) with the following conditions: mobile phase A (chloroform:methanol:ammonium hydroxide, 89.5:10:0.5) and mobile phase B (chloroform:methanol:ammonium hydroxide:water, 55:39:0.5:5.5). MRM transitions were set up for comparative analysis of various polar lipids. Individual lipid species were quantified by referencing to spiked internal standards, including d_31_-PC(16:0/18:1), d_31_-PE(16:0/18:1), d_31_-PG(16:0/18:1), d_31_-PS(16:0/18:1), d_7_-PI(15:0/18:1), PA 17:0/17:0, CL 80:4, LPC-d_4_-26:0, LPE 17:1, LPI 17:1 and LPS 17:1 from Avanti Polar Lipids. Free fatty acids were quantitated using d_31_-16:0 (Sigma-Aldrich) and d_8_-20:4 (Cayman Chemicals) as internal standards. Glycerol lipids including DAG and TAG were quantified using a modified version of reverse phase HPLC/MRM. Separation of neutral lipids was achieved on a Phenomenex Kinetex-C18 2.6 μm column (i.d. 4.6×100 mm) using an isocratic mobile phase containing chloroform:methanol:0.1 M ammonium acetate 100:100:4 (v/v/v) at a flow rate of 300 μL for 10 min. Levels of short-, medium-, and long-chain TAGs were calculated by referencing to spiked internal standards of TAG(14:0)_3_-d_5_, TAG(16:0)_3_-d_5_ and TAG(18:0)_3_-d_5_ obtained from CDN isotopes. DAGs were quantified using d_5_-DAG16:0/16:0 and d_5_-DAG18:1/18:1 as internal standards (Avanti Polar Lipids). To facilitate comparison between samples, the lipid content was normalized to total protein in each sample.

### Statistical analysis

All statistical analysis was done in GraphPad Prism 8.0 (GraphPad Software, Inc.). Each point stands for one independent experimental repeat and all error bars indicate SEM. Data was first inspected for normality. Nonparametric tests were applied when the data deviated from normal distribution. Otherwise, the two-tailed unpaired t-test was used for comparison between two groups; the test was with Welch’s correction if the groups had unequal SD. Ordinary one-way analysis of variance (ANOVA) was used for comparison among multiple groups with equal SD, and Brown-Forsythe and Welch ANOVA was used for multiple groups with unequal SD followed by an appropriate *post hoc* test for multiple comparisons (for each experiment, the tests used are stated in the figure legend). Statistically significant differences between groups with two variations were analyzed by two-way ANOVA. Statistical significances are indicated as: * *P* < 0.05, ** *P* < 0.01, *** *P* < 0.001, **** *P* < 0.0001.

## Acknowledgements

We thank Dr. Jingyan Zhang, Dr. Pingsheng Liu and Dr. Xiaochen Wang for providing reagents and helpful discussions. We thank Andi Liu and Kang Xie for initial observations. This research was supported by the National Natural Science Foundation of China and the Ministry of Science and Technology of China (9195420001 and 2018YFA0506902).

## Author contributions

X.H. supervised the study. J.Z. and X.H. wrote the manuscript. J.Z. performed the most of experiments and analysis. S.M.L. and G.S. extracted and measured the lipids. X.H. and L.Y. performed the EMS screen. J.L. and M.D. performed the EM microscopy. The authors declare they have no competing interests.

## Data and materials availability

All data needed to evaluate the conclusions in the paper are present in the paper and/or the Supplementary Materials. Additional data as well as materials used in this study may be requested from authors.

## Notes

### Competing Interest Statement

The authors have declared no competing interest.

