## Supplemental Figure S1, S2 and Supplemental Table 1 for "Contrasting effects of reduced phosphatidylcholine synthesis in absence of seipin: embryonic lethality goes down while lipid droplets get even larger"

### Supplementary Materials

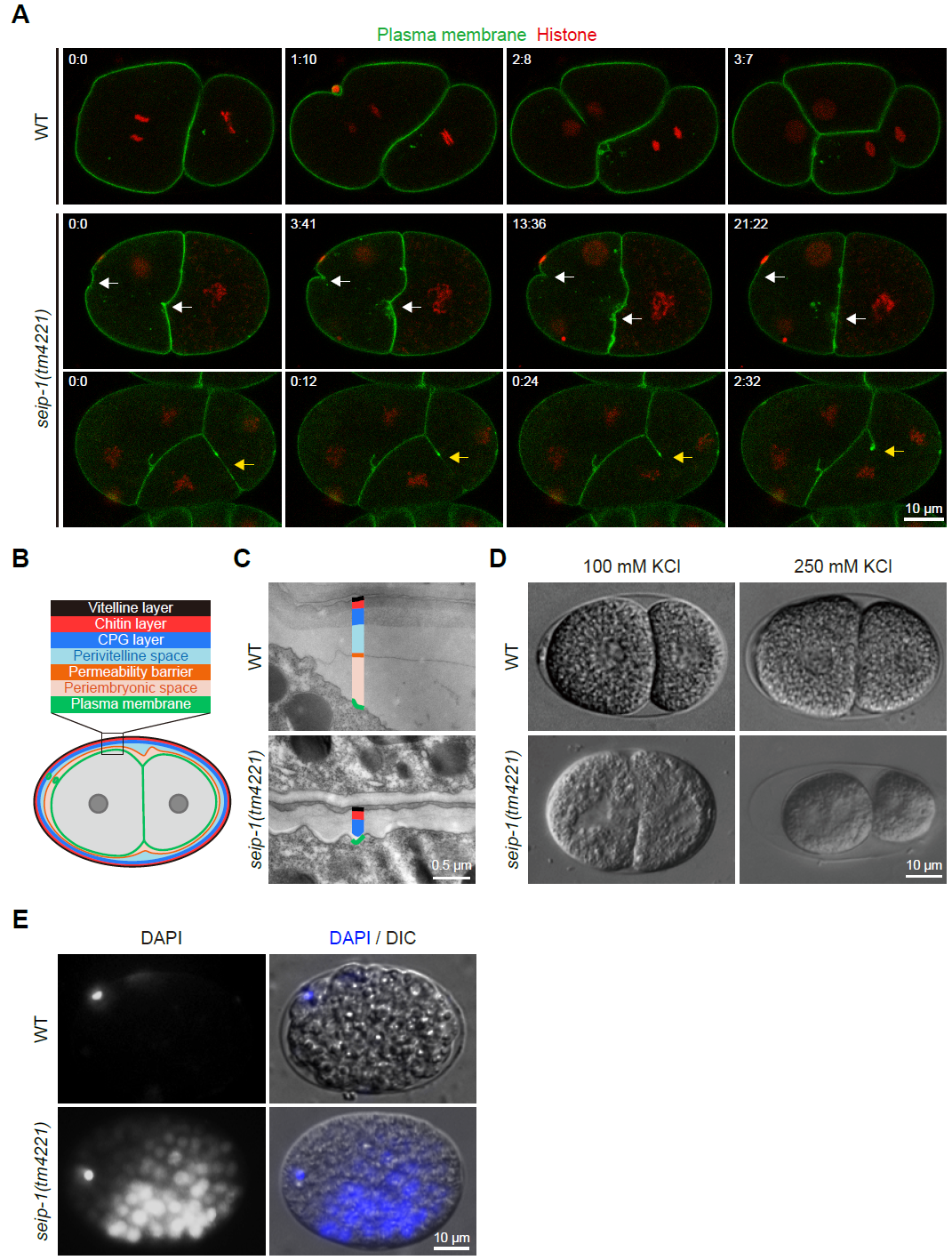

**Fig. S1. *seip-1(tm4221)* mutants have permeable eggshells.**

(A) Time-lapse confocal images of wild-type and *seip-1(tm4221)* embryos at the early cleavage stage. Plasma membranes were labelled by GFP::PH(PLC1delta1) and nuclei were labelled by mCherry::Histone. The relative time points are shown at the top left. White arrows indicate that cytokinesis proceeds more slowly and aborts halfway. Yellow arrows indicate cell fusion mediated by abrupt disruption of the adjoining plasma membrane.

(B) Diagram of the composition of the *C. elegans* eggshell. Six eggshell layers are distinguished by different colors outside the plasma membrane. The first and second polar bodies are located in the perivitelline space and periembryonic space, respectively.

(C) Transmission electron microscopy images of wild-type and *seip-1(tm4221)* eggshells. The eggshell layers are distinguished by colors according to B.

(D) Differential interference contrast microscopy (DIC) images of wild-type and *seip-1(tm4221)* embryos which were soaked in hypotonic (100 mM KCl) or hypertonic (250 mM KCl) solution.

(E) Fluorescence and DIC images of wild-type and *seip-1(tm4221)* embryos which were stained with DAPI.

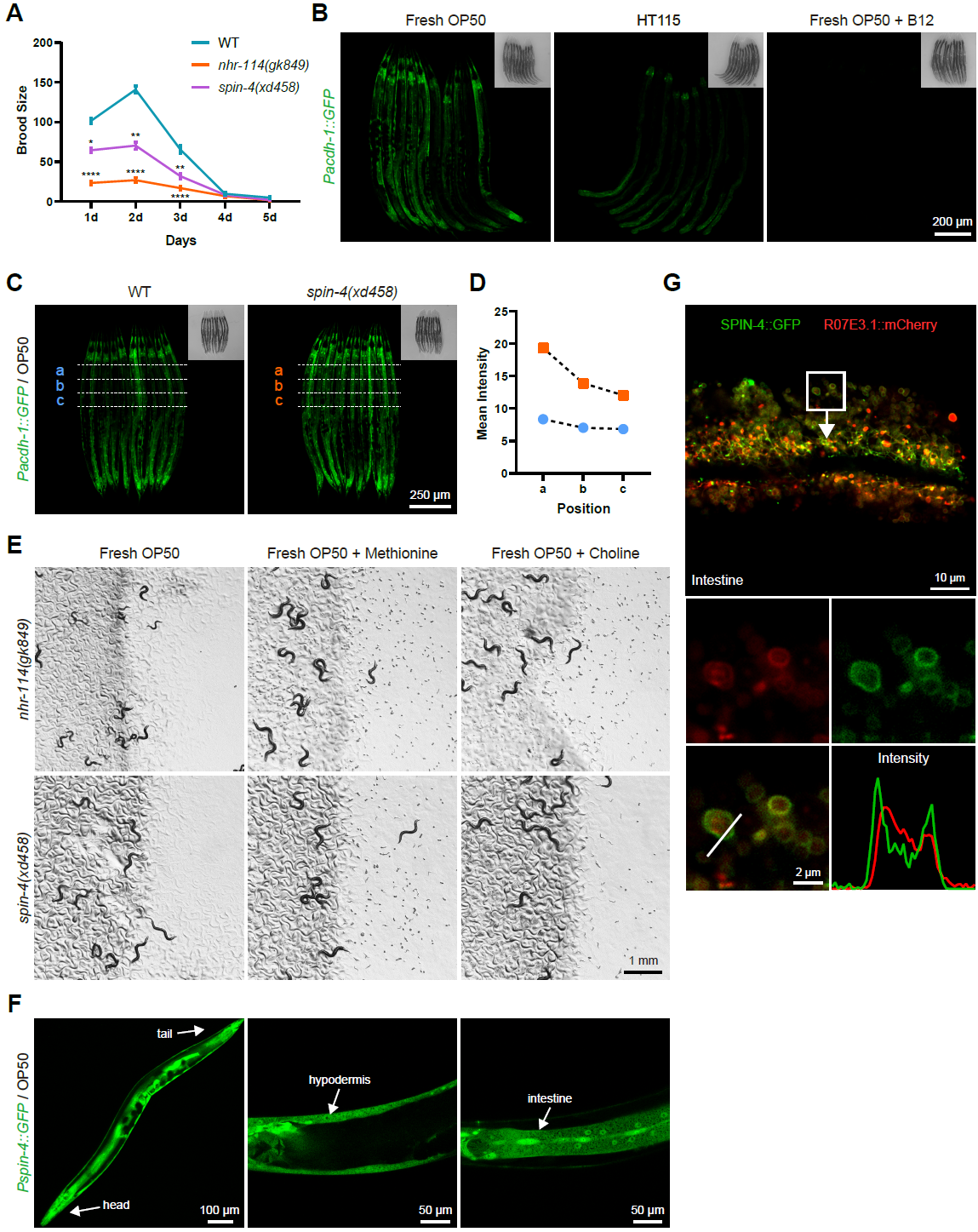

**Fig. S2. The sterility of *nhr-114* and *spin-4* mutants is diet-dependent.**

(A) Quantification of brood size of wild type, *nhr-114(gk849)* mutants and *spin-4(xd458)* mutants over five time periods from the beginning of adulthood. Error bars represent SEM. The numbers of tested animals for wild type, *nhr-114(gk849)* and *spin-4(xd458)* are 13, 20 and 18, respectively. Statistically significant differences between wild type and each mutant at each time point were determined by nonparametric test (Kruskal-Wallis H-test) with *post hoc* Dunn’s test: **P* < 0.05, ***P* < 0.01, *****P* < 0.0001.

(B) Confocal images of *Pacdh-1::GFP* reporter animals fed on different diets with or without B12 supplement. Insets are bright-field images.

(C) Confocal images of the *Pacdh-1::GFP* reporter in the wild-type or *spin-4(xd458)* background. Insets are bright-field images. Three regions of each image were segmented to calculate the mean fluorescence intensity.

(D) Quantification of fluorescence intensity at the three different positions in C.

(E) Bright-field images of worm plates carrying *nhr-114(gk849)* or *spin-4(xd458)* mutants on fresh OP50 supplied with or without 10 mM methionine or 50 mM choline.

(F) Confocal images of adult *Pspin-4::GFP* animals. The fluorescence of *Pspin-4::GFP* is observed throughout the worm, including head, tail, hypodermis and intestine, as indicated by the white arrows.

(G) Confocal images of adult animals expressing SPIN-4::GFP and mCherry::R07E3.1. The enlarged images of the boxed area show that the fluorescence of SPIN-4::GFP encircles the signal from the mCherry-tagged lysosomal peptidase R07E3.1. The line profile of the signals is shown at the bottom right.

**Table S1. *C. elegans* strains used in this study.**

| **Strain** | **Source** | **Genotype** |
| --- | --- | --- |
| N2 | CGC | wild type (Bristol) |
| CB4856 | CGC | wild type (Hawaiian) |
| XD4613 | NBRP(Japan) | *seip-1(tm4221) V* |
| XD3546 | Huang Lab | *seip-1(tm4221) V; xdEx1640 [Pseip-1::seip-1 + rol-6(su1006)]* |
| RT130 | CGC | *pwIs23 [vit-2::GFP]* |
| XD5440 | Huang Lab | *seip-1(tm4221) V; pwIs23 [vit-2::GFP]* |
| JIM113 | CGC | *unc-119(ed3) III; ujIs113 [Ppie-1::mCherry::H2B + Pnhr-2::mCherry::HIS-24 + unc-119(+)] II* |
| OD58 | CGC | *unc-119(ed3) ltIs38 [Ppie-1::GFP::PH(PLC1delta1) + unc-119(+)] III* |
| XD4901 | Huang Lab | *ujIs113 [Ppie-1::mCherry::H2B + Pnhr-2::mCherry::HIS-24 + unc-119(+)] II; ltIs38 [Ppie-1::GFP::PH(PLC1delta1) + unc-119(+)] III* |
| XD4866 | Huang Lab | *seip-1(tm4221) V; ujIs113 [Ppie-1::mCherry::H2B + Pnhr-2::mCherry::HIS-24 + unc-119(+)] II; ltIs38 [Ppie-1::GFP::PH(PLC1delta1) + unc-119(+)] III* |
| XD1017 | Huang Lab | *seip-1(tm4221) III/nT1 [qIs51] (IV;V)* |
| XD3714 | Huang Lab | *seip-1(tm4221) V; spin-4(xd286) III* |
| XD3715 | Huang Lab | *seip-1(tm4221) nhr-114(xd287) V* |
| XD5260 | Huang Lab | *spin-4(xd458) III* |
| XD5303 | Huang Lab | *seip-1(tm4221) V; spin-4(xd458) ltIs38 [Ppie-1::GFP::PH(PLC1delta1) + unc-119(+)] III* |
| XD4860 | Huang Lab | *seip-1(tm4221) nhr-114(xd428) V* |
| XD6124 | Huang Lab | *seip-1(tm4221) nhr-114(xd428) V; ltIs38 [Ppie-1::GFP::PH(PLC1delta1) + unc-119(+)] III* |
| VC1760 | CGC | *nhr-114(gk849) V* |
| XD5905 | Huang Lab | *spin-4(xd458) ltIs38 [Ppie-1::GFP::PH(PLC1delta1) + unc-119(+)] III* |
| XD6122 | Huang Lab | *nhr-114(gk849) V; ltIs38 [Ppie-1::GFP::PH(PLC1delta1) + unc-119(+)] III* |
| RB1434 | CGC | *mmcm-1(ok1637) III* |
| RB755 | CGC | *metr-1(ok521) II* |
| QC122 | CGC | *paqr-2(tm3410) III; pcyt-1(et9) X* |
| XD6503 | Huang Lab | *pcyt-1(et9) X; ujIs113 [Ppie-1::mCherry::H2B + Pnhr-2::mCherry::HIS-24 + unc-119(+)] II* |
| VL749 | CGC | *wwIs24 [Pacdh-1::GFP + unc-119(+)]* |
| XD6333 | Huang Lab | *spin-4(xd458) III; wwIs24 [Pacdh-1::GFP + unc-119(+)]* |
| XD5403 | Huang Lab | *xdEx2409 [Pspin-4::GFP + Podr-1::RFP]* |
| XW7951 | Xiaochen Wang | *qxIs448 [R07E3.1::mCherry]* |
| XD5553 | Huang Lab | *qxIs448 [R07E3.1::mCherry]; xdEx2483 [Pvha-6::SPIN-4::GFP + Podr-1::RFP]* |
| XD6500 | Huang Lab | *seip-1(tm4221) V; mmcm-1(ok1637) III* |
| XD6502 | Huang Lab | *seip-1(tm4221) V; metr-1(ok521) II* |
| XD6504 | Huang Lab | *seip-1(tm4221) V; pcyt-1(et9) X; ujIs113 [Ppie-1::mCherry::H2B + Pnhr-2::mCherry::HIS-24 + unc-119(+)] II* |
| XD6339 | Huang Lab | *seip-1(tm4221) V; wwIs24 [Pacdh-1::GFP + unc-119(+)]* |
| BX30 | CGC | *fat-3(wa22) IV* |
| XD6294 | Huang Lab | *seip-1(tm4221) V; fat-3(wa22) IV; ltIs38 [Ppie-1::GFP::PH(PLC1delta1) + unc-119(+)] III* |
| XD6295 | Huang Lab | *seip-1(tm4221) V; spin-4(xd458) III; fat-3(wa22) IV* |
| XD6296 | Huang Lab | *seip-1(tm4221) nhr-114(xd428) V; fat-3(wa22) IV; ltIs38 [Ppie-1::GFP::PH(PLC1delta1) + unc-119(+)] III* |
